# Atmospheric plasma jet device for versatile electron microscope grid treatment

**DOI:** 10.1101/2021.05.11.443639

**Authors:** Eungjin Ahn, Tianyu Tang, Byungchul Kim, Hae June Lee, Uhn-Soo Cho

## Abstract

Atmospheric pressure plasmas have been widely applied in surface modification and biomedical treatment due to its ability to generate highly reactive radicals and charged particles. In negative-stain electron microscopy (Neg-EM) and cryogenic electron microscopy (cryo-EM), plasmas have been used in eliminating the surface contaminants as well as generating the hydrophilic surface to embed the specimen on grids. Plasma treatment is a prerequisite for negative stain and quantifoil grids, which are coated with hydrophobic carbon on the grid surface. Here we introduce a non-thermal atmospheric plasma jet system as an alternative new tool for surface treatment. Unlike the conventional glow discharger, we found that the plasma jet system successfully cleans the grid surface and introduces hydrophilicity on grids in the ambient environment without introducing a vacuum. Therefore, we anticipate the plasma jet system will be beneficial in many aspects, such as cost-effective, convenient, versatile, and potential applications in surface modification for both negative stain and cryo-EM grid treatment.

## Introduction

Solving atomic-resolution protein structure using single-particle cryogenic electron microscopy (cryo-EM) becomes more and more routine and substantial in the structural biology field. Recent technical advances in cryo-EM instruments (cold field-emission gun, aberration corrector, next-generation direct detector, etc.), as well as software developments, further accelerate this trend (1, 2). Therefore, it is not surprising to see that sample and grid preparation become a bottleneck in illustrating the protein structure using cryo-EM. Purified biomolecular assemblies are embedded in thin amorphous ice after plunge-frozen in liquid ethane. Successful microscopic data acquisition requires the optimization of specimen preparation procedures, such as selecting grid types/treatment and finding the best blotting condition (3, 4). Various types of grids are commercially available and have been examined to overcome ice thickness, particle drafting during image collection (5), protein denaturation due to the exposure in the air-water interface (6–8). Regardless of grid types, plasma treatment is a prerequisite before applying protein specimens (9). Since the grid surface is hydrophobic and contaminated by dirt and others, plasma treatment is a necessary step to clean and modify the grid surface into hydrophilic, thereby enhancing grid and solution contact. Particularly, negative stain EM grids and the quantifoil grids are additionally coated with hydrophobic carbons, which must be modified to the hydrophilic surface before applying the specimen (10).

Plasma treatment for material processing has a long history and successful achievement in the semiconductor industry. For the last couple of decades, low-temperature plasmas (LTPs) in atmospheric pressure have been applied to an enormous range of biomedical applications and surface treatment by virtue of the controllability of the chemical reactions of radicals and the change in the energy distributions of charged particles (11–13). LTPs have non-equilibrium electrons and ions where the gas temperature and the ion temperature are at the range of room temperature not to deliver thermal damage on the body, while the electron temperature is up to tens of thousands Kelvin (14). High energy electrons generate complex chemical species and high ion flux by the collisions with neutral gas. Ions can be accelerated across thin sheath layers to impact the surface (15). The synergy effect of the radicals and the ions on the surface is well understood for the semiconductor etching process in low-pressure plasma processing (16), but it is much more complicated to understand the interaction of atmospheric pressure plasmas because of high collisionality.

The low-energy plasma modification system is a widely used instrument for grid pre-treatment to introduce hydrophilicity and clean the grid surface under the vacuum. Recently, the plasma treatment step combined with vapors of chemical precursors has been used to further introduce functional groups on the graphene surface to overcome preferred orientation and reduce specimen movement (17). Although this new approach looks promising in functionalizing the graphene surface in a controlled manner, it is very challenging to build such a device as an individual laboratory. Here, we introduce an atmospheric pressure plasma jet device that generates plasma in the air environment with the potential of versatile surface modification. The strengths of the plasma jet system are: First, it is easy to install at a low price. Second, it is easy to use without introducing a vacuum. Finally, with the assistance of additional setups, it has the potential for surface modification with simple chemical molecules. We anticipate that this plasma jet device can greatly contribute to the development of functional grids and single particle protein observation in cryo-EM.

## Result

### Building atomospheric pressure plasma jet device

Plasma is described as a quasi-neutral mixture of charged particles and radicals in a partially ionized gas. Those activated gas molecules possess high reactivity toward the hydrocarbon surface, which changes their chemical state into oxidized forms and makes them hydrophilic for further applications. In general, surface treatment by plasma requires a vacuum environment to increase the carry distance (*i.e.*, mean free path) of activated ionic gas molecules, otherwise, it can be eliminated before arriving at the target surface due to its high collision rate and reactivity with other gas molecules. The mean free path of an ionized gas molecule is calculated by using Equation 1, where *λ_i_* is the mean free path of the gas molecule *i*, *k* is the Boltzmann constant (1.38 × 10^−23^ J/K), *T* and p are the temperature and pressure of the environment, respectively, and d_i_ is the diameter of a gas molecule *i*:

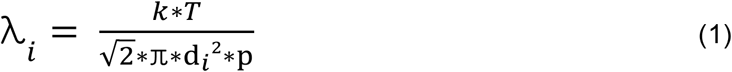

For example, the mean free path of activated Argon (Ar) molecules at the ambient condition (298 K, 1 atm) is shorter than 70 nm, while it could be increased up to 300 μm in a vacuum environment (298 K, 0.20 mbar). The plasma jet system helps charged particles to reach the target surface by pumping with a carrier gas before making an unfavorable reaction in the atmosphere, thereby proceeding surface modification even in the atmospheric environment. The plasma jet device composition only requires several parts such as a DC voltage source, a high voltage DC-AC inverter circuit, a dielectric tube (*i.e.*, jet part), and gas flow controlled with the mass flow controller (Fig. 1a). Therefore, the plasma jet system does not require a vacuum pump or a chamber and works well in the atmospheric environment, which also helps reducing operation time without any warm-up. The setup price for the whole plasma jet device, including AC circuit, power source, carrier gas, and other safety tools, is less than $ 700 and could be set up easily in a day without any professional knowledge or experience.

**Figure 1.**
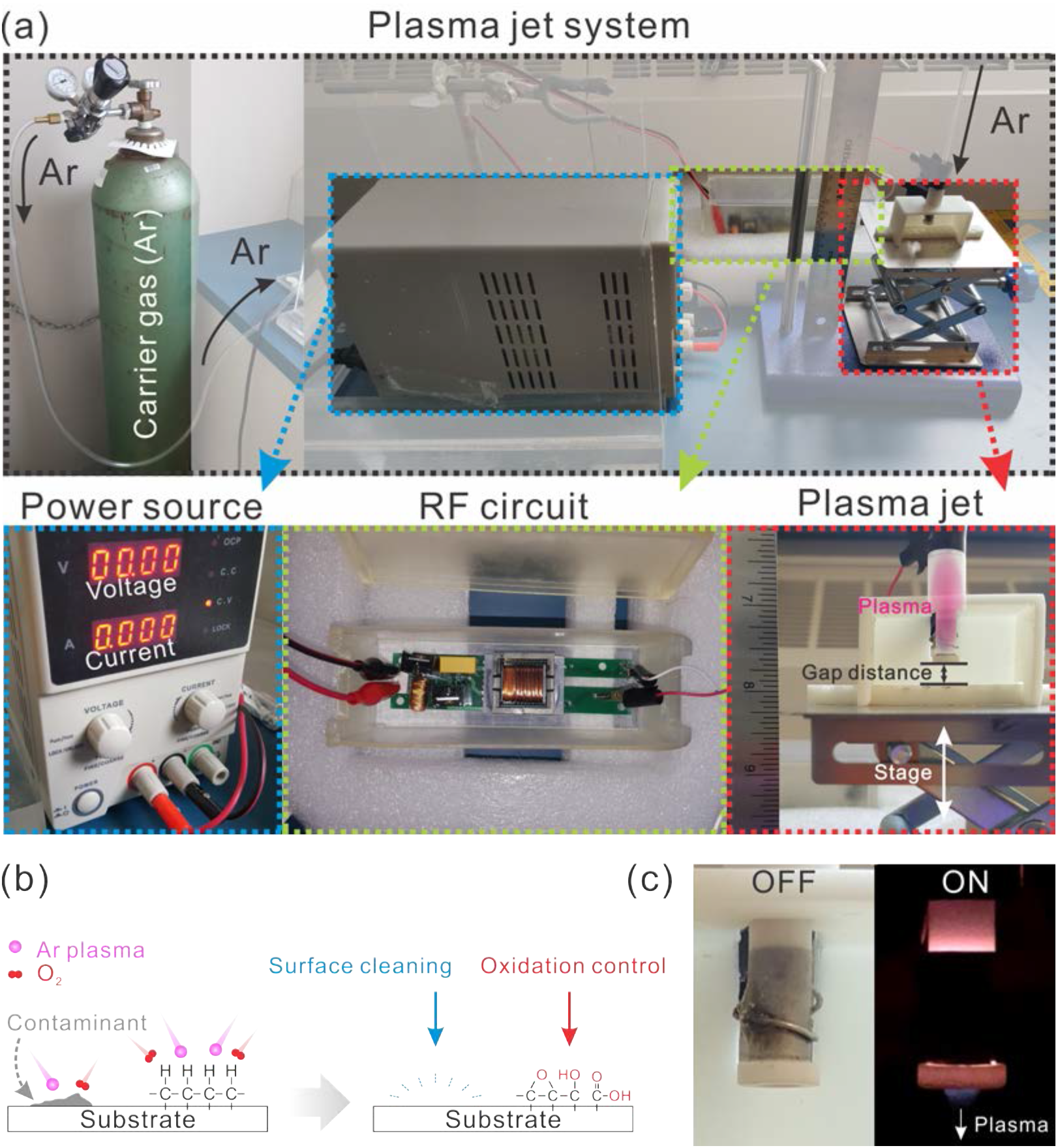
(a) Photograph of the overall plasma jet system (black dotted box) with each component highlighted by dotted box: power source (blue), high-voltage DC-AC inverter circuit (green), jet part (red). (b) Schematic illustration of the plasma treatment effect on the hydrocarbon surface (c) Photograph of the dielectric jet part (left) and plasma ejected from the jet under the dark background (right).

Typical chemical reactions in the plasma jet tube occur as follows. First, carrier gas (Ar) is activated into either cationic or radical (excited species are purple colored for clarification) forms via the high voltage generated by the AC circuit with the following chemical reaction R2 and R3 (18):

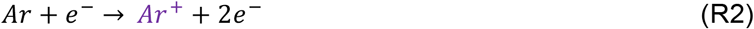

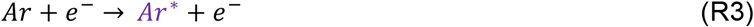

Then those activated Ar species have larger energy (> 11.5 eV) than the ionization threshold of O_2_ molecule (11.09 eV) which generate the activated oxygen species (R4 and R5) (19, 20), otherwise the excess energy is released in the form of light (R6):

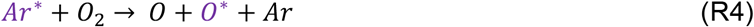

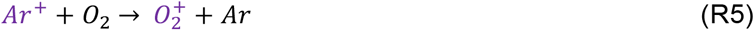

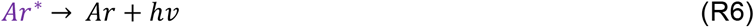

When the excited oxygen species reach the hydrocarbon surface contaminated by dirts and others, gradual oxidation occurs and removes those contaminants. It also generates functional groups (*e.g.*, hydroxyl, epoxy, carboxyl), which converts the hydrocarbon surface to the polar and hydrophilic surface (Fig. 1b). The plasma jet is activated when a minimum power (0.55 W) is applied to the AC circuit with a gas flow injected from the tube to the surface (Fig. 1c). The detailed inner structure of the jet part and applied voltage versus discharge current waveforms of the Ar plasma jet is described in Fig. S1. The discharge power consumed by the Ar plasma jet under this condition was calculated as 0.43 W by the Lissajous figure method with a 0.1 μF external capacitor.

### Hydrophilicity and surface cleaning effect mediated by the Plasma Jet

The hydrophilicity and surface cleaning effect was compared between the plasma jet and commercial glow discharger (PELCO easiGlow, vacuum pressure < 0.26 mbar). After plasma jet treatment on the petri dish, the water droplet spread along the surface due to increased hydrophilicity (Fig. 2a). To monitor the degree of surface modification upon the plasma jet treatment, the water contact angle goniometer was employed. The gap distance between the jet part and specimen was altered along with various input power (Fig. S2). Hydrophilicity on the petri dish increases along with shorter gap distance and higher input power (Fig. 2b). Not always the lower contact angle (*i.e.*, high hydrophilicity) surfaces enhance the biocompatibility with proteins (21). Therefore, we decided to set the gap distance of 1 cm and the power of 1.07 W for the plasma jet treatment because this condition can induce a moderate level of hydrophilicity on the target surface, which is compatible with the commercial glow discharger.

**Figure 2.**
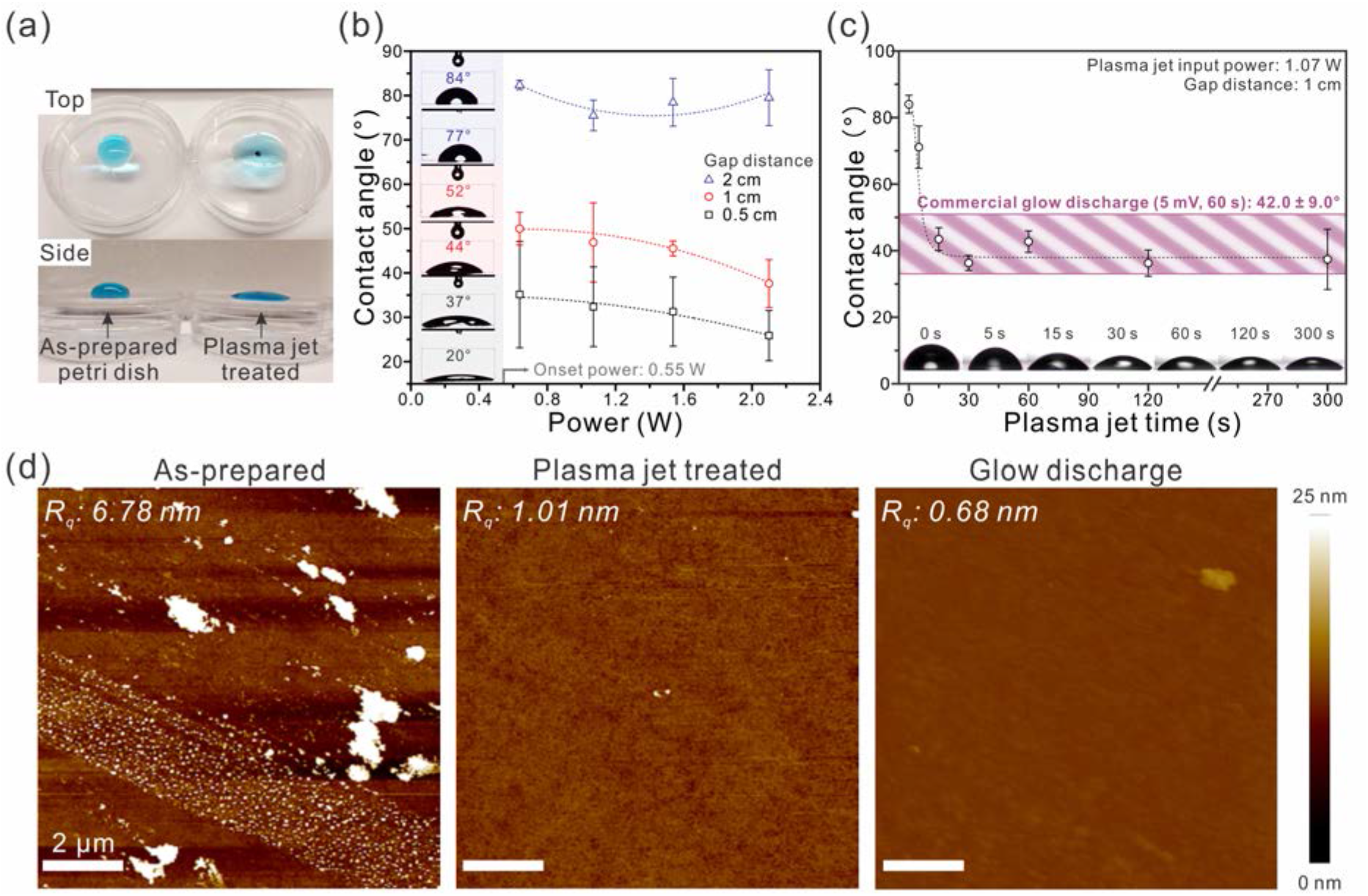
(a) Hydrophilicity change on the petri dish before and after plasma jet treatment (2 min). The water droplet was colored with methylene blue to help visualization. (b, c) Water contact angle monitored by the contact angle goniometer on the petri dish surface after plasma jet treatments in (b) different input powers and gap distances and (c) time-dependent contact angle in fixed input power (1.07 W) and gap distance (1 cm). The violet box area denotes the average contact angle of petri dish treated by commercial glow discharge (5 mV, 60 s). Each plot in contact angle goniometer results was averaged after 5-time measurements in different spots. Insets: Photographes of water droplet with the measured contact angle and plasma jet treated time. (d) AFM morphology comparison before (as-prepared) and after plasma jet treatment and glow discharge treated slide glass.

Time-dependent plasma jet treatment with the fixed gap distance (1 cm) and input power (1.07 W) showed that the contact angle of the plasma jet treated surface over 15 seconds was comparable to measured the contact angle of glow discharge treated surface (5 mA, 60 s), which was 42 ± 9.0° (Fig. 2c) (22). The atomic force microscopy (AFM) was introduced to evaluate the surface cleaning effect after plasma jet treatment. We compared the surface roughness and morphology on a glass slide before and after plasma jet and glow discharge treatment (Fig. 2d). The surface roughness (R_q_) was 6.78 nm in as-prepared slide glass, while it was reduced to 1.01 nm and 0.68 nm after plasma jet and glow discharge treatment, respectively. This denotes that the plasma jet system is effective in surface cleaning and inducing hydrophilicity and is comparable to the commercial glow discharge device without introducing a vacuum. When the plasma jet was applied to the negative stain grid, it was shown that the water droplet shape was spread evenly through the surface of the grid, compared to poor wetting before plasma treatment (Fig. S3). This implies that the protein samples could be uniformly coated on the surface without aggregations. This plasma-treated grid was used for the next step, protein observation by the negative stain method.

### Oxidation studies of plasma treatment with graphene

Detailed oxidative behavior of plasma jet device was studied by using graphene as a substrate, as it is easy to understand the oxidation state of monolayerd graphene by using Raman spectroscopy. To compare the oxidation power of the plasma jet system, we used the commercial graphene monolayer coated on the Cu foil (Graphenea) and monitored the degree and type of oxidation upon time-dependent plasma treatment. The plasma jet treatment showed a comparable oxidation level to the glow discharger, with gradually increasing D peak (1340 cm^−1^) intensity by time, which indicates the defect density accumulation upon exposure (Fig. 3a, b). Both conditions similarly maintained the G peak (1580 cm^−1^), which denotes the maintained basal plane of graphene. However, a notable difference between the plasma jet and glow discharger happened at the 2D peak (2670 cm^−1^) when we treat the plasma over 2 min. The graphene treated by 5 min glow discharge showed a decreased and broadened 2D peak intensity, which indicates a diminish of graphitic ordered regions in the graphene lattice (23, 24) compared to the plasma jet treated graphene. Therefore, Raman spectra analysis demonstrated that the plasma jet system has the potential to modify the surface of the graphene monolayer to introduce oxidation. Moreover, the plasma jet system showed a better effect in maintaining the graphene lattice (2D peak) compared to the commercial glow discharge.

**Figure 3.**
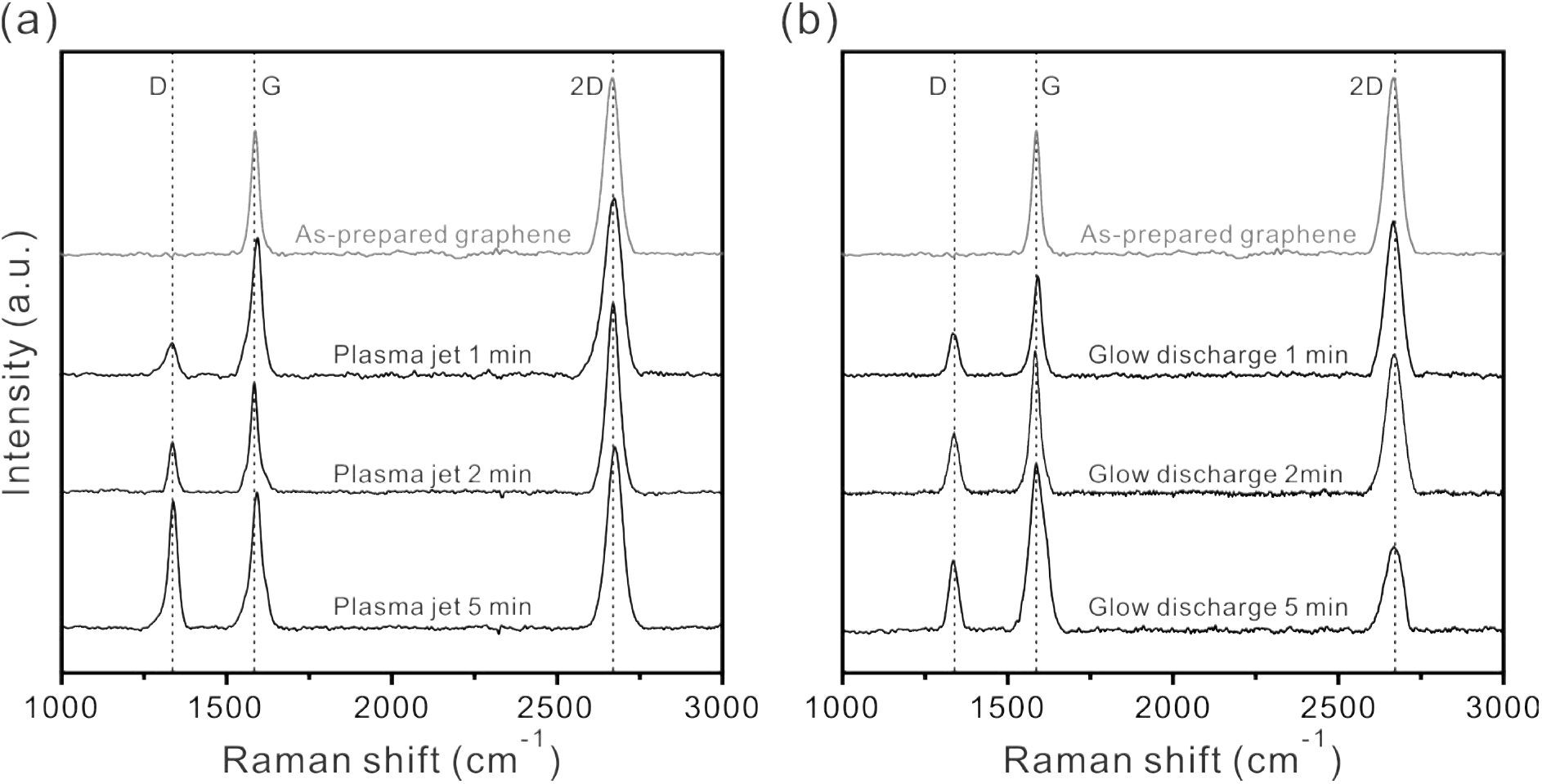
Raman spectra of the graphene monolayers on the Cu foil treated with (a) the plasma jet and (b) the glow discharge. Three vertical dotted lines indicated the position of D, G, and 2D peaks from the graphene lattice, respectively.

### Comparison of negative-stain EM images using the conventional glow discharger and the plasma jet system

To monitor the practical application of the plasma jet system on EM grids, we employed the negative-stain EM approach. The negative-stain EM grids are coated with amorphous carbons, which maintain the hydrophobic surface. The plasma treatment introduces hydrogenation on carbon-coated EM grids, thereby allowing the solution specimen to make direct contact with the grid surface. Three different grids–untreated, plasma jet treated (1.07 W, 60 s), and glow discharger treated (5 mA, 60 s)–were prepared and used to apply the protein sample (*Methylococcus capsulatus* soluble methane monooxygenase, *M. caps* MMOH) using the standard protocol. As shown in Fig. 4 and Fig. S4, the plasma jet treated negative-stain EM grid showed well-distributed particles on the surface comparable to the one treated with the glow discharger, unlike the untreated negative-stain EM grid, which showed particle/stain aggregation on the surface. This indicates that the plasma jet device can successfully introduce hydrophilicity and cleaning on the grid surface comparable to the commercial glow discharger.

**Figure 4.**
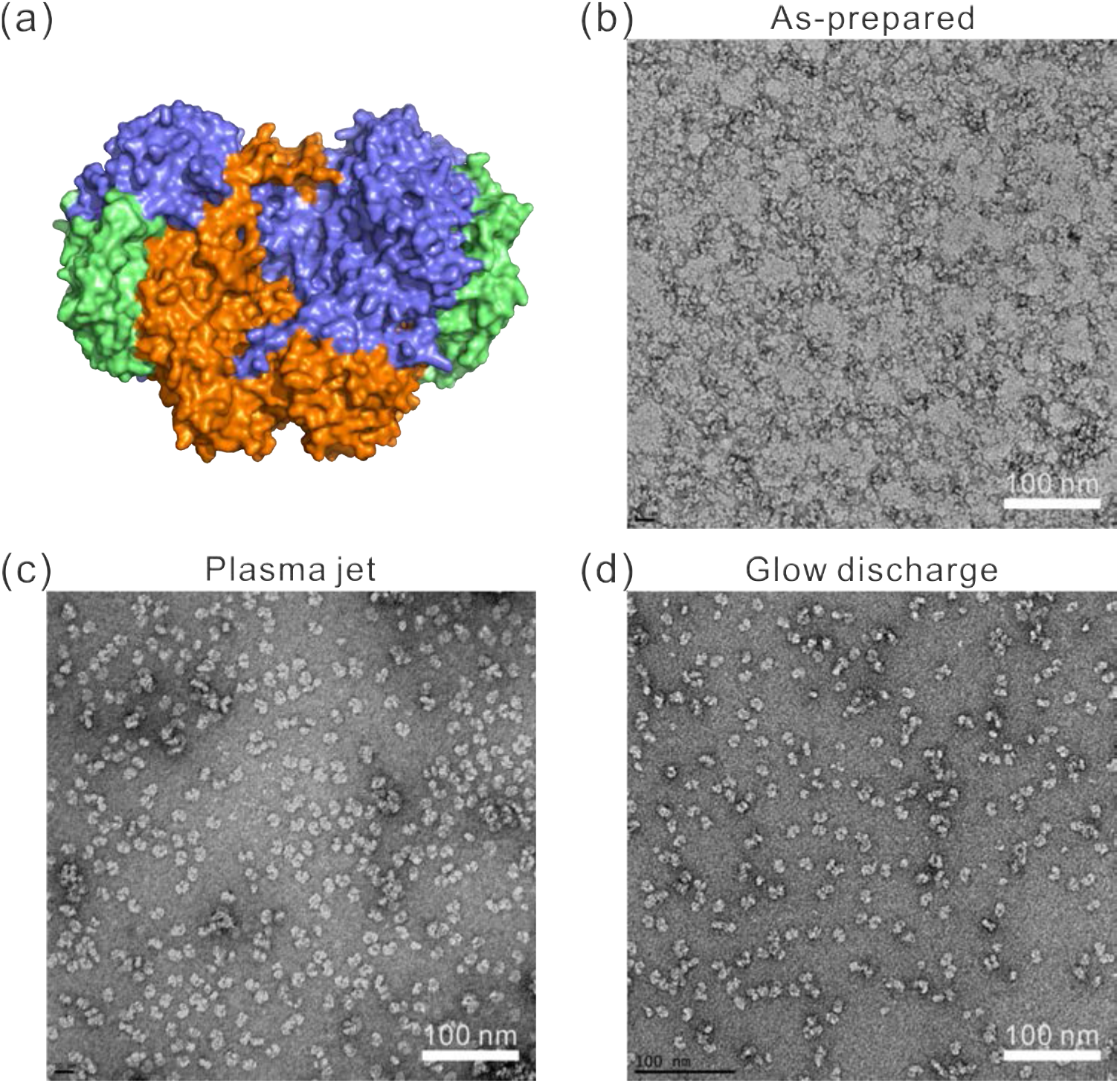
(a) Surface representation of *M. caps* MMOH and typical negative-stain images of MMOH on (b) as-prepared, (c) plasma jet, and (d) glow discharge-treated negative-stain grids.

### Cryo-EM analysis using the plasma jet treated grid

Plasma treatment on the cryo-EM grid is a prerequisite step for specimen application and the following vitrification. To examine whether the plasma jet system is further applicable for cryo-EM grid preparation, we treated the gold (Au) quantifoil grid (Ted Pella, Inc) using the plasma jet (1.07 W, 60 s) and prepared the cryo-EM grid using *M. caps* MMOH as a test specimen. Blotting (5 sec) and plunge-freezing on liquid ethane were performed using the Vitrobot (Thermo-Fisher), and data screening/collection was done using 200 KeV Arctica with a K2 summit direct detector. Obtained microscopic images indicate that particles are well dispersed with no obvious preferred orientation (Fig. 5a and S5). Particles showed enough contrast clearly visible under the defocus ranges of −1.0 ~ −2.0 μm, which suggests that ice-thickness is well controlled in plasma jet treated cryo-EM grids. Total 310 microscopic images were collected for particle picking and data quality analysis. 2D classes obtained from ~38k particles visualized the clear secondary structures (Fig. 5b), and we could obtain a subnanometer resolution (6.4 Å) structure after the 3D reconstruction (Fig. 5c). Therefore, the structural study of *M. caps* MMOH indicated that the plasma jet is suitable for cryo-EM analysis.

**Figure 5.**
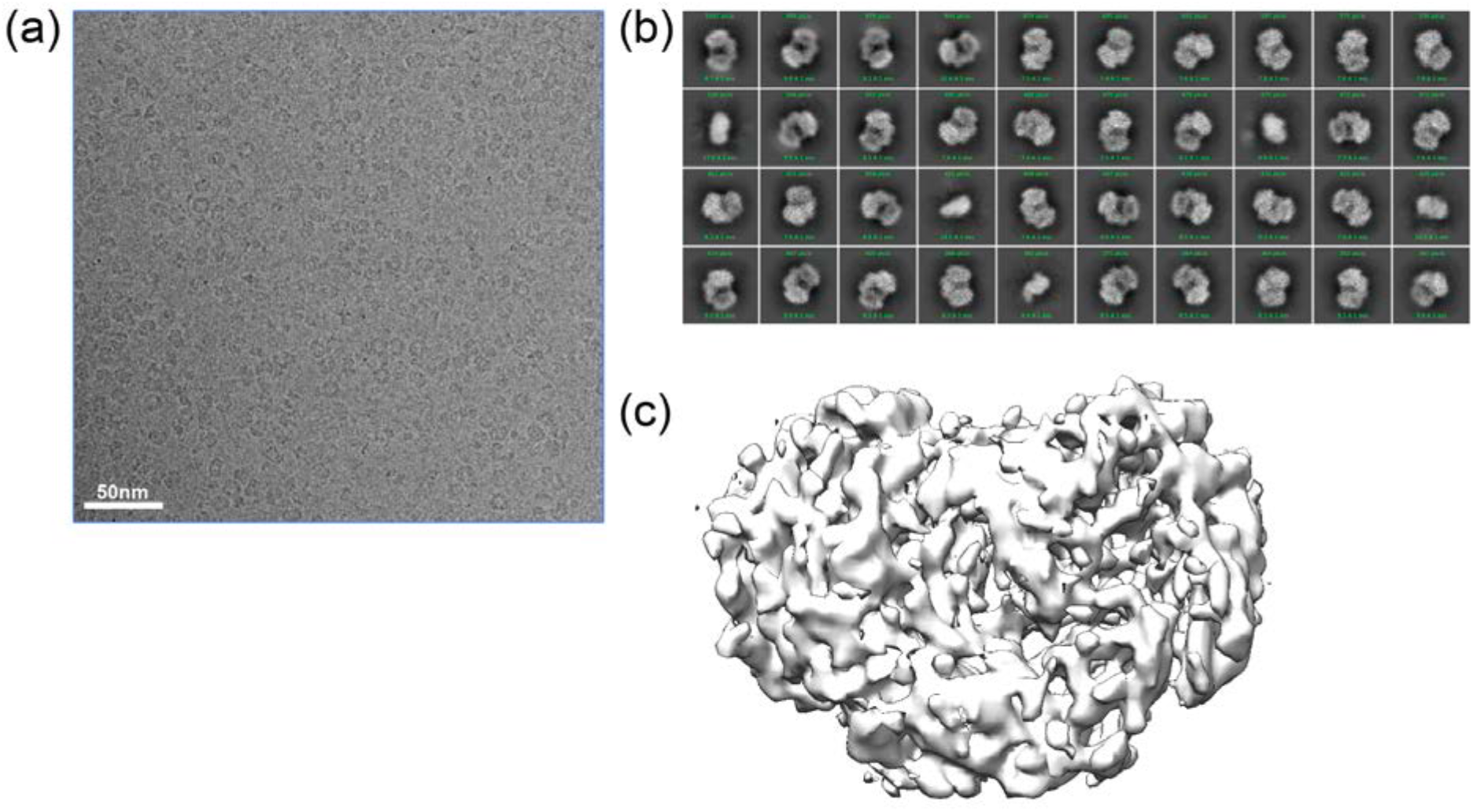
Cryo-EM analysis of *M. caps* MMOH using the plasma jet treated Au quantifoil grid. (a) A representative microscopic image of *M. caps* MMOH. (b) Top 40 classes of the 2D classification (out of 100 classes). (c) The 3D reconstructed structure of *M. caps* MMOH at 6.4 Å resolution.

In this study, we describe the atmospheric plasma jet system that can be utilized as a surface polishing and hydrophilic treatment tool on both Neg-EM and cryo-EM grids. The plasma jet system has strength in cost-effective, easy to set up, available with versatile use, and most importantly, it operates in the atmospheric environment without the vacuum system. We demonstrated that the plasma jet system has the potential to replace the conventional glow discharge method by comparing their effectiveness in hydrophilicity, surface cleaning, and oxidations with water contact angle goniometer, AFM, Raman studies.

### Experimental procedures

#### Atmospheric plasma jet setup

We utilize a plasma jet, which is a dielectric barrier discharge with the guided gas flow in a tube. A cylindrical metal rod with a radius of 1.5 mm was set at the center of the ceramic tube with an inner radius of 2 mm. A layer of copper tape with a width of 5 mm was covered at the outer ring of the alumina ceramic tube. The thickness of the dielectric material (ceramic tube) is 1 mm. Plasma is generated between the metal rod and the ceramic tube, a ring area with an inner radius of 1.5 mm and an outer radius of 2 mm, and diffuses out through the nozzle. The inner electrode was connected to a high-voltage power source which generates a sinusoidal waveform voltage at 1-3 kV with a frequency of 20 kHz.

#### Contact angle goniometer

A contact angle goniometer (Ossila) was used to measure the water contact angle before and after the plasma-treated surface. The water droplet was dropped on the target surface, and a polynomial curve is fitted to the droplet edge with the contact angle above 10**°**. Where the curve crosses the surface baseline, it’s tangent line is used to determine the contact angle.

#### Raman spectroscopy

Raman microscopy (Renishaw) system equipped with a 532 nm diode laser and a 1200 lines/mm grating was used for spectrum collection through an Olympus SLMPlan 20× objective. All spectra were obtained in extended scan mode in the range of 3000-1000 cm^−1^ for analysis of framework bands, peak position from 2680 cm^−1^ for analysis of the 2D band, 1580 cm^−1^ and 1380 cm^−1^ for monitoring the G and D band of graphene, respectively. Calibration of the laser was performed in static scan mode using a silicon standard.

#### Atomic force microscope (AFM)

AFM images were taken using a Veeco Dimension Icon Atomic Force Microscope with a ScanAsyst-Air AFM tip from Bruker Nano Inc. The data were analyzed using Nanoscope Analysis 2.0 software.

#### Negative-stain electron microscopy (Neg-EM)

Carbon supported negative-stain EM grids were prepared either without (as-prepared) or with following plasma treatment. The plasma jet grid was treated with applied power of 1 W at the atmosphere and the commercial glow discharge grid was treated with 5 mA for 1 min under vacuum (< 26 mbar). *M. caps* MMOH, purified as previously described (25), was immobilized on one of these grids followed by addition of uranyl formate for enhancing contrast and carrying out negative-stain EM. The negative-stain EM micrographic images were collected using 100 kV Morgagni (FEI) at the University of Michigan cryo-EM center.

#### Cryogenic electron microscopy (cryo-EM)

The 300 mesh Au R1.2/1.3 quantifoil grid (Electron Microscopy Sciences) was pretreated with the plasma jet (1.07 W, 60 s). 5 μl of *M. caps* MMOH (0.1 mg/ml) was applied to the Au quantifoil grid and plunge-frozen using a Mark IV Vitrobot (Thermo Fisher Scientific) with 5 sec blotting time. Grids were mounted on 200 KeV Talos Arctica (Thermo Fisher Scientific) and microscopic images were taken 45,000 X in a counting mode with the pixel size of 0.98 Å/pixel at liquid nitrogen temperature. A dose rate of 1.52 electrons/Å^2^/frame and defocus values ranging from −1.0 to −2.0 μm were used. Total exposure of 7 sec per image was dose-fractionated into 40 movie frames, resulting in an accumulated dose of 64.11 electrons per Å^2^. A total of 310 movies were collected and particle picking was done using the Topaz program (26) embedded in the cryoSPARC software package (27). Total 66,482 particles were picked and following 2D/3D classification were done using the cryoSPARC software package. The final reconstructed 3D structure (6.4 Å resolution) was generated from 23,403 particles (Fig. 5c).

## Abbreviations

The abbreviation used are:

Neg-EM: negative-stain electron microscopy
cryo-EM: cryogenic electron microscopy;
LTP: low-temperature plasmas
DC: direct current
AC: alternating current
*M. caps* MMOH: *Methylococcus caps* soluble methane monooxygenase
AFM: atomic force microscope
SLM: standard liter per min

## Acknowledgments

We thank Dr. Leila Foroughi and Dr. Adam Matzeger in the Department of Chemistry (the University of Michigan) for the use of the Raman spectroscopy instrument and assistance. We thank staffs at the University of Michigan cryo-EM center for the use of the Morgagni instrument and assistance. We thank Dr. Haiping Sun and the Michigan Center for Materials Characterization for the use of the AFM instrument and assistance. Dr. Seung-Jae Lee at the Chonbuk National University kindly provided the purified *M. caps* MMOH. This research was supported by NIH DK111465 / CA250329 to U.-S.C. and PNU-RENovation (2020-2021) to H.L.

## Author contributions

U.-S. C. and H. L. conceived the project. E. A. carried out all the experiments. T. T. helped setting up the plasma jet system for the lab scale. B. K. performed a negative stain EM. E. A. and U.-S. C. wrote the manuscript.

## Conflicts of interests

The authors declare that they have no conflicts of interest with the contents of this article.

## Notes

### Competing Interest Statement

The authors have declared no competing interest.

### Summary of Updates

New data was included

