## Supplemental figure for "Atmospheric plasma jet device for versatile electron microscope grid treatment"

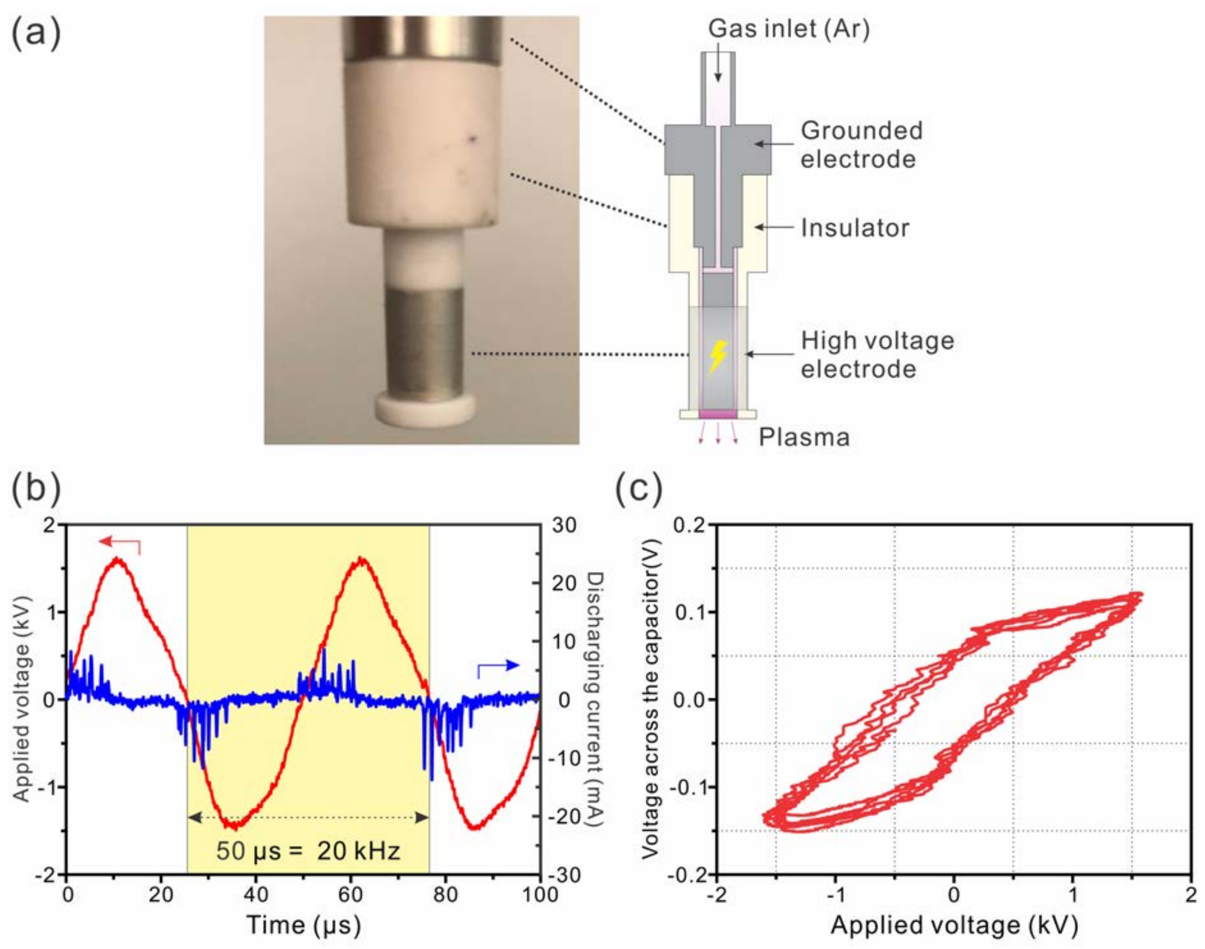


**Figure S1**. (a) Photograph and illustration of the inner components of the Ar plasma jet device (b) Waveforms of the applied voltage and the discharge current and (c) Lissajous figure of the Ar plasma jet. Applied peak-to-peak voltage and the gas flow rate are 3.2 kV and 2 SLM.

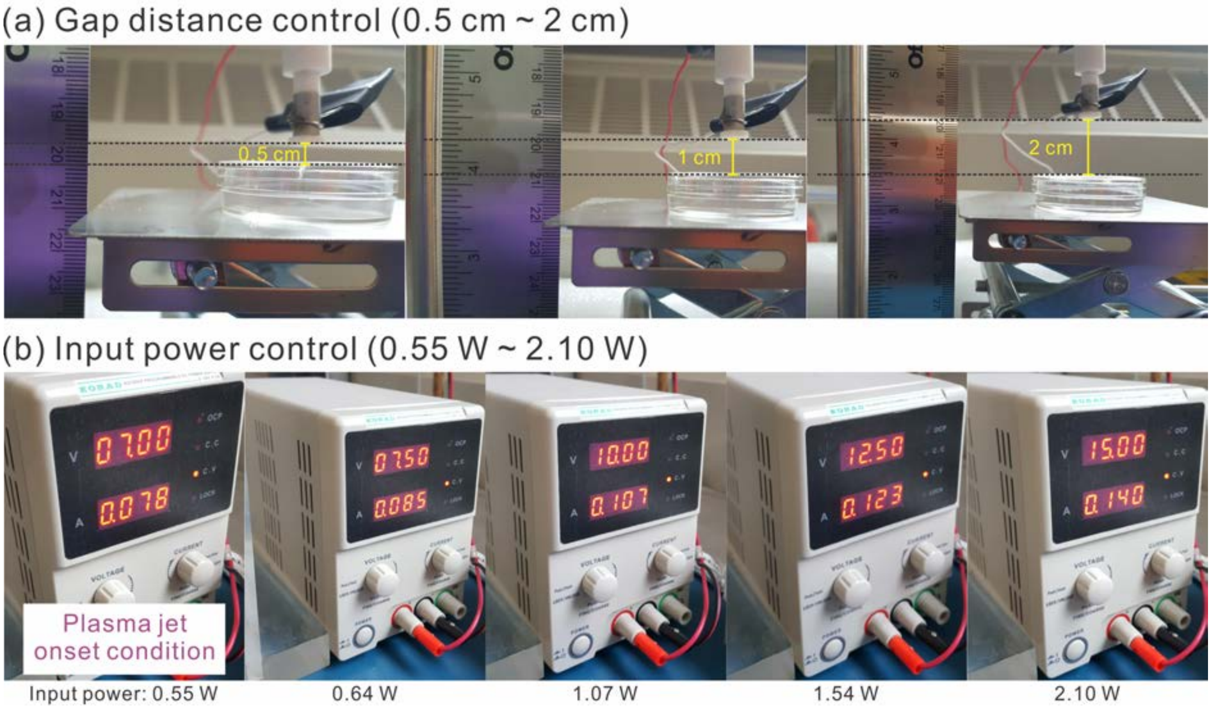


**Figure S2**. Photograph of plasma jet conditions on petri dish with different (a) gap distances (0.5 cm ~ 2 cm) and (b) input powers (0.55 W ~ 2.10 W).

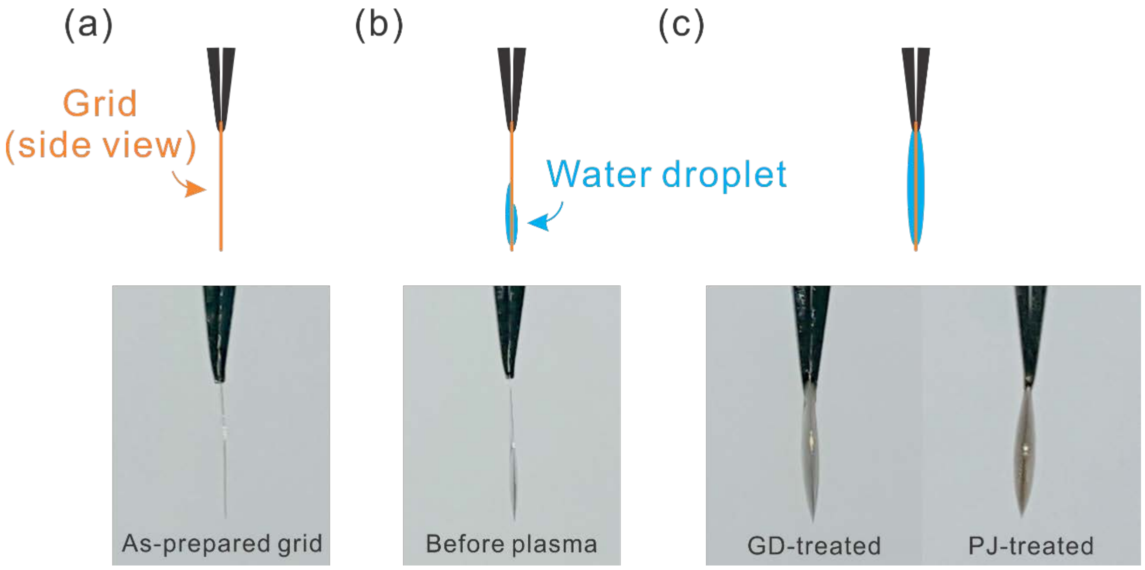


**Figure S3**. (a) Schematic side view and photograph of a negative-stain EM grid. Wettability test with DI water (b) before and (c) after glow discharge (GD) and plasma jet (PJ) treatment.

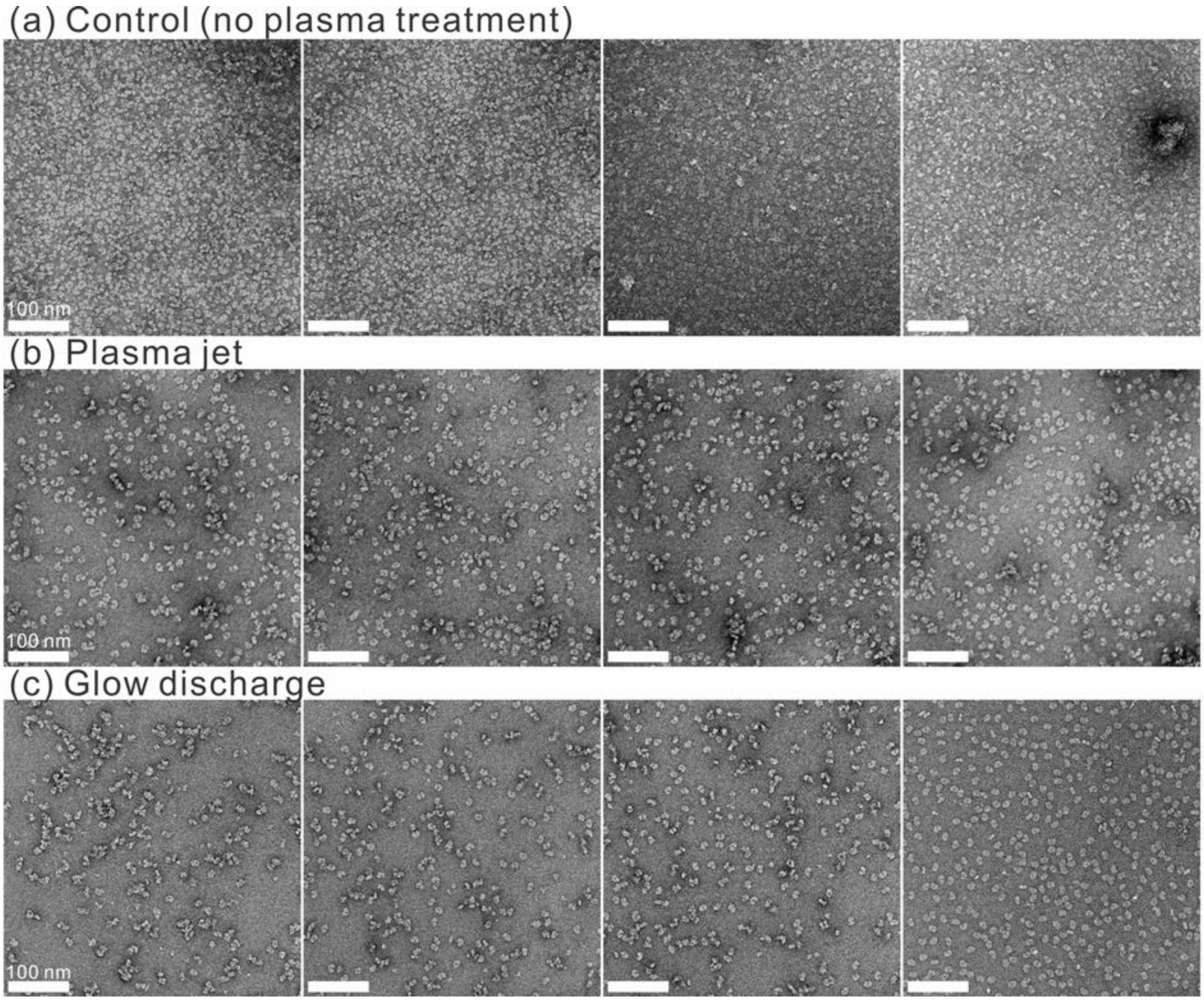


**Figure S4**. Negative-stain images of MMOH on (a) as-prepared, (b) plasma jet, and (c) glow discharge-treated negative-stain grids.

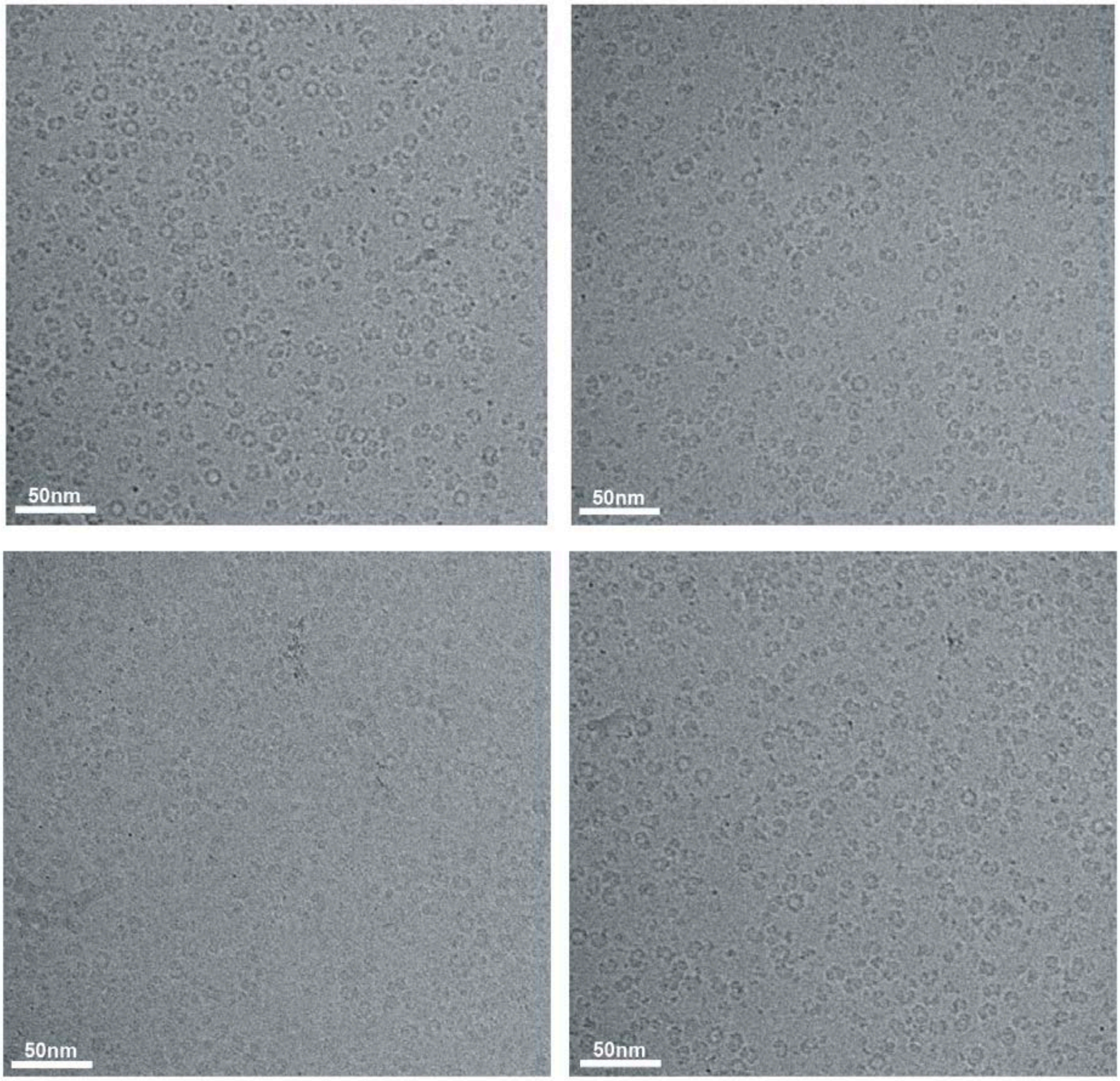


**Figure S5**. Additional cryo-EM microscopic images of *M. caps* MMOH from the plasma jet-treated Au quantifoil grid.
